# Distinct functional domains of Dystroglycan regulate inhibitory synapse formation and maintenance in cerebellar Purkinje cells

**DOI:** 10.1101/2024.08.29.610348

**Authors:** Jennifer N. Jahncke, Eric Schnell, Kevin M. Wright

## Abstract

Dystroglycan is a cell adhesion molecule that localizes to synapses throughout the nervous system. While Dystroglycan is required to maintain inhibitory synapses from cerebellar molecular layer interneurons (MLIs) onto Purkinje cells (PCs) whether initial synaptogenesis during development is dependent on Dystroglycan has not been examined. We show that conditional deletion of *Dystroglycan* from Purkinje cells prior to synaptogenesis results in impaired MLI:PC synapse formation and function due to reduced presynaptic inputs and abnormal postsynaptic GABA_A_ receptor clustering. Using genetic manipulations that disrupt glycosylation of Dystroglycan or truncate its cytoplasmic domain, we show that Dystroglycan’s role in synapse function requires both extracellular and intracellular interactions, whereas synapse formation requires only extracellular interactions. Together, these findings provide molecular insight into the mechanism of inhibitory synapse formation and maintenance in cerebellar cortex.

## Introduction

Throughout nervous system development, synapses are formed, refined, and eliminated. Synapse formation begins when a presynaptic axon recognizes potential postsynaptic sites through molecular cues. After initial contact between the pre- and postsynaptic neurons is established, transmembrane and secreted molecules are recruited to the nascent synapse to regulate its maturation and refinement in response to activity (Südhof, 2018). The wide diversity in cell adhesion molecules (CAMs) present at synapses implies that they can form a “code” for determining synapse type and specificity. While several CAMs have been shown to be involved in the maturation and maintenance of synapses, less is known about the molecular players involved in initial synaptogenesis.

Dystroglycan is a CAM expressed throughout the nervous system in a variety of cellular populations including neurons, astrocytes, oligodendrocytes, vascular endothelial cells, and neuroepithelial cells (Colognato et al., 2007; Nguyen et al., 2014; Nickolls and Bönnemann, 2018; Tian et al., 1996; Zaccaria et al., 2001). Dystroglycan is encoded by a single gene (*Dag1*) that undergoes post-translational autoproteolytic cleavage to produce two subunits: α- and β-Dystroglycan (Holt et al., 2000; Ibraghimov-Beskrovnaya et al., 1992). The extracellular alpha subunit is extensively glycosylated through a process involving at least 19 different genes, and employs a specific glycan motif referred to as “matriglycan” to bind several extracellular proteins that contain Laminin-G (LG) domains (Jahncke and Wright, 2023). Mutations that disrupt α-Dystroglycan glycosylation lead to a form of congenital muscular dystrophy referred to as dystroglycanopathy that is frequently accompanied by a wide range of neurodevelopmental abnormalities (Nickolls and Bönnemann, 2018). The transmembrane beta subunit of Dystroglycan noncovalently binds the alpha subunit in the extracellular environment, and contains a short C-terminal intracellular domain that binds multiple scaffolding and signaling proteins (Moore and Winder, 2010). Dystroglycan is the transmembrane component of the Dystrophin Glycoprotein Complex (DGC), which functions to connect the actin cytoskeleton to the extracellular matrix. Dystroglycan and Dystrophin, which interacts with Dystroglycan’s intracellular domain, are both central to the DGC, whereas other components of the complex can vary depending on the cellular type.

Early in nervous system development, Dystroglycan plays important roles in neuronal migration, axon targeting, and maintenance of the blood-brain barrier (Clements et al., 2017; Clements and Wright, 2018; Lindenmaier et al., 2019; Menezes et al., 2014; Myshrall et al., 2012; Wright et al., 2012). Later in development, as synapses begin to form, Dystroglycan expression increases in several neuronal populations throughout the brain, where it localizes at inhibitory synapses (Briatore et al., 2020, 2010; Brünig et al., 2002; Lévi et al., 2002; Patrizi et al., 2008). Dystroglycan is required for the formation and function of subsets of inhibitory synapses onto hippocampal and cortical pyramidal neurons, where it appears to function as a postsynaptic recognition cue for presynaptic CCK^+^/CB_1_R^+^ basket interneuron axons (Früh et al., 2016; Jahncke et al., 2024; Miller and Wright, 2021). Dystroglycan expression continues past the period of synapse formation in pyramidal neurons, and is required to maintain CCK^+^/CB_1_R^+^ basket synapses (Früh et al., 2016). However, whether Dystroglycan plays these same roles at other synapses has not been explored in detail. In the cerebellum, Dystroglycan is present at inhibitory somatic and dendritic synapses onto Purkinje cells and deletion of *Dag1* from Purkinje cells after synaptogenesis results in a gradual reduction of inhibitory synapse number and impaired synaptic function (Briatore et al., 2020, 2010; Patrizi et al., 2008). Constitutive loss of *Dystrophin* results in similar alterations in inhibitory inputs onto Purkinje cells, confirming an important role for the DGC at these synapses (Kueh et al., 2008; Wu et al., 2022). However, the molecular mechanisms by which Dystroglycan regulates inhibitory synapses are unclear.

Until recently, tools for the conditional deletion of genes in cerebellar Purkinje cells prior to the period of synaptogenesis have been lacking. We identified *Calb1^Cre^*as a line that drives recombination in embryonic Purkinje cells, allowing us to conditionally delete *Dag1* from Purkinje cells prior to the onset of synapse formation (Jahncke and Wright, 2024). Using multiple deletion strategies, we show that Dystroglycan is required for both inhibitory synapse formation and maintenance in Purkinje neurons. Furthermore, we provide *in vivo* mechanistic insight by showing that extracellular glycosylation of α-Dystroglycan and intracellular interactions through β-Dystroglycan play distinct roles in synapse formation and function.

## Results

### Dystroglycan is exclusively localized to inhibitory synapses in cerebellar cortex

Dystroglycan is associated with populations of inhibitory synapses throughout the brain (Briatore et al., 2010, 2020; Jahncke et al., 2024; Trotter et al., 2023; Früh et al., 2016). To rigorously assess Dystroglycan’s synaptic localization in cerebellar cortex, we conducted analysis of Dystroglycan immunoreactivity co-localization with established markers of synaptic populations. Inhibitory synapses in cerebellar cortex reflect synapses from (1) molecular layer interneurons (MLIs) onto Purkinje cells (PCs), (2) MLIs onto other MLIs, and (3) PC collaterals onto other PCs. While currently available markers are unable to distinguish between these three populations of synapses, Dystroglycan-positive synapses are presumed to be from MLIs onto PCs (Briatore et al., 2010) and potentially reflect a population that is at least partially distinct from the Gephyrin-containing population of synapses (Lévi et al., 2002; Uezu et al., 2019). We therefore used two presynaptic inhibitory synapse markers (VGAT and GAD67) to analyze inhibitory MLI:PC synapses. To analyze MLI:PC synapses specifically, we masked the synaptic channels using the Purkinje cell marker Calbindin, eliminating any immunofluorescent signal outside of Purkinje cells. The IIH6 antibody, which specifically recognizes matriglycan disaccharide repeats on α-Dystroglycan, significantly co-localized with both VGAT and GAD67, with nearly 50% of IIH6 puncta positive for VGAT or GAD67 (Error! Reference source not found. **A**, **C**). As a control, we flipped the synaptic marker channel along the vertical axis (VGAT or GAD67) to generate a mirror image of the original channel and calculated co-localization between the original IIH6 channel and the mirrored synaptic marker channel. IIH6 co-localized with the original channel significantly more than the mirrored channel for both VGAT and GAD67 (Error! Reference source not found. **A-D**), confirming that Dystroglycan is localized to inhibitory synapses in cerebellar cortex.

There are two populations of excitatory synapses onto Purkinje cells: (1) VGluT1^+^ parallel fiber inputs from granule cells within the cerebellar cortex and (2) VGluT2^+^ climbing fiber inputs originating from neurons of the inferior olivary nucleus. Despite the high density of VGluT1^+^ parallel fibers, Dystroglycan and VGluT1 immunoreactivity were mutually exclusive such that mirroring the VGluT1 channel increased the incidence of Dystroglycan:VGluT1 overlap (**Supplemental Figure 1 A-B**). Co-localization between IIH6 and VGluT2 was minimal, and was not affected by mirroring the VGluT2 channel, indicating that Dystroglycan does not localize to climbing fiber synapses (**Supplemental Figure 1 C-D**). Together, this illustrates that Dystroglycan expression is exclusively localized to inhibitory synapses in cerebellar cortex.

### Dystroglycan is required for the formation and function of inhibitory synapses onto Purkinje cells

While recent work has shown that Dystroglycan is required at CCK^+^/CB_1_R^+^ basket interneuron synapses onto pyramidal neurons in the hippocampus and cortex, our understanding of Dystroglycan’s role at inhibitory cerebellar synapses remains incomplete due to the limited genetic tools available to manipulate cell populations prior to synapse formation in the cerebellum (Briatore et al., 2020; Früh et al., 2016; Jahncke et al., 2024). We recently identified *Calb1^Cre^* as a tool for conditional deletion of *Dag1* from embryonic Purkinje cells, allowing us to investigate Dystroglycan’s role in the initial formation of inhibitory synapses onto Purkinje cells (Jahncke and Wright, 2024). We crossed *Calb1^Cre^;Dag1^+/-^*mice with a *Dystroglycan* conditional line (*Dag1^flox/flox^*) to generate *Calb1^Cre^;Dag1^flox/-^* conditional knockouts (*Calb1^Cre^;Dag1^cKO^*) and littermate *Calb1^Cre^;Dag1^flox/+^* controls (*Calb1^Cre^;Dag1^Ctrl^*). Due to the restricted expression of *Cre* to Purkinje cells in the cerebellum, this approach allowed us to investigate the role of postsynaptic Dystroglycan at MLI:PC synapses in a cell-autonomous manner.

In order to assess inhibitory synapse function in *Calb1^Cre^;Dag1^Ctrl^*and *Calb1^Cre^;Dag1^cKO^* cerebella, we recorded mini inhibitory postsynaptic currents (mIPSCs) from Purkinje cells at P25. On average, *Calb1^Cre^;Dag1^cKO^* Purkinje cells exhibited reduced mIPSC amplitude and frequency compared to littermate controls (**Figure 2 A-F**). Reduced mIPSC frequency is generally an indicator of a reduction in the number of synapses, whereas reduced amplitude could reflect either (1) changes to postsynaptic receptor clustering resulting in disrupted subsynaptic domains or (2) altered subcellular distribution of synapses driven by a reduction in the number of postsynaptic GABA_A_ receptors. There was no difference in either mIPSC rise or decay time between control and conditional knockout Purkinje cells (Error! Reference source not found.), suggesting no substantial changes in the subunit composition of GABA_A_ receptors between genotypes.

**Figure 1.**
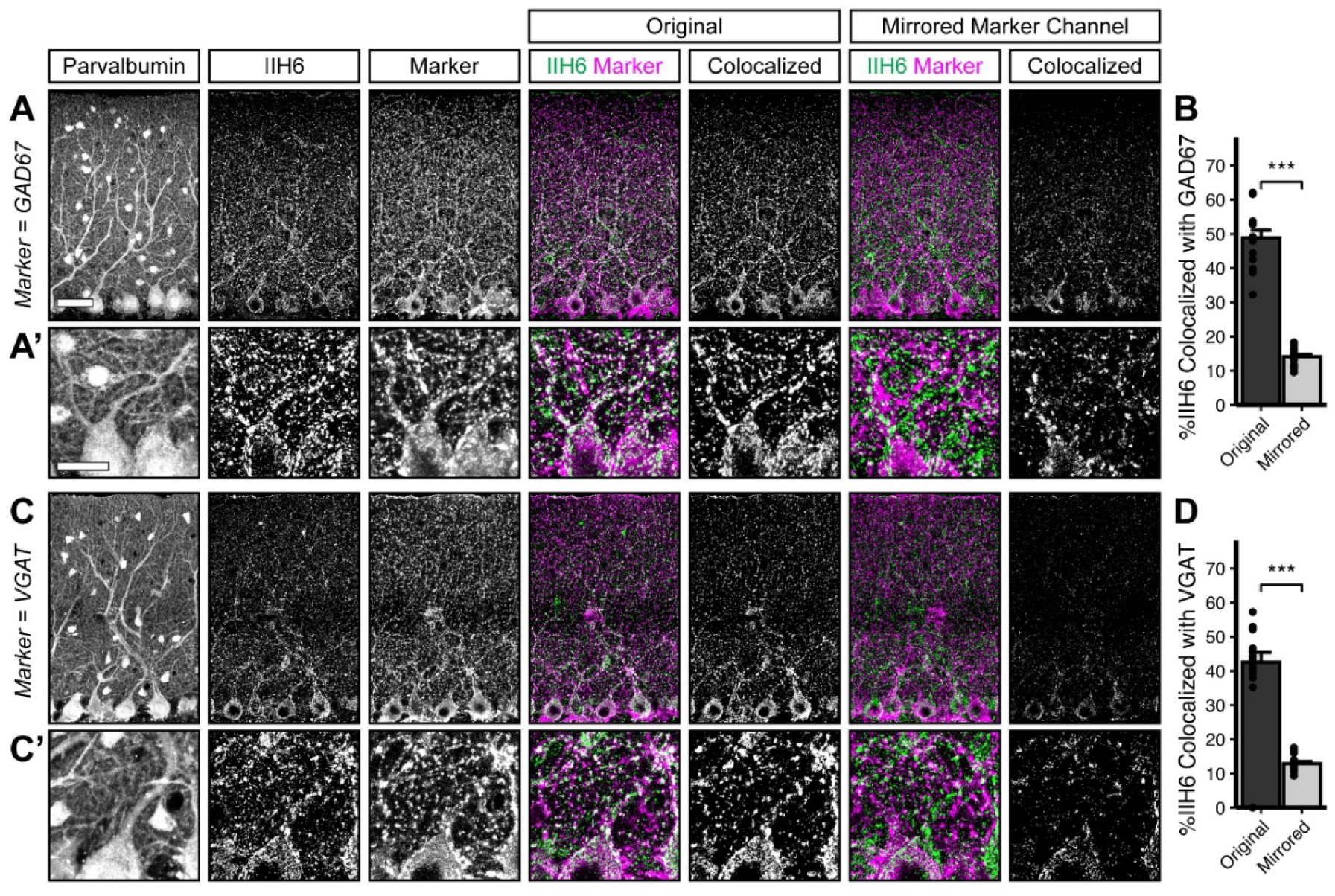
Dystroglycan co-localizes with markers of inhibitory synapses. Cerebellar cortex of lobules V-VI were immunostained with Parvalbumin to show Purkinje cell and MLI morphology and counterstained with IIH6 (glycosylated Dystroglycan) and GAD67 (**A**) or VGAT (**C**). Both the merged channels (IIH6, green; GAD67/VGAT, magenta) and colocalized pixels are shown for the original image and for original IIH6 with the mirrored GAD67/VGAT channel. Images are maximum projections. (**B, D**) Quantification of the percent of IIH6 puncta that are colocalized with GAD67/VGAT puncta. Scale bar for (**A**, **C**) is 50μm; scale bar for insets (**A’**, **C’**) is 25μm. GAD67 N = 16 images, 3 animals. VGAT N = 18 images, 4 animals.

**Figure 2.**
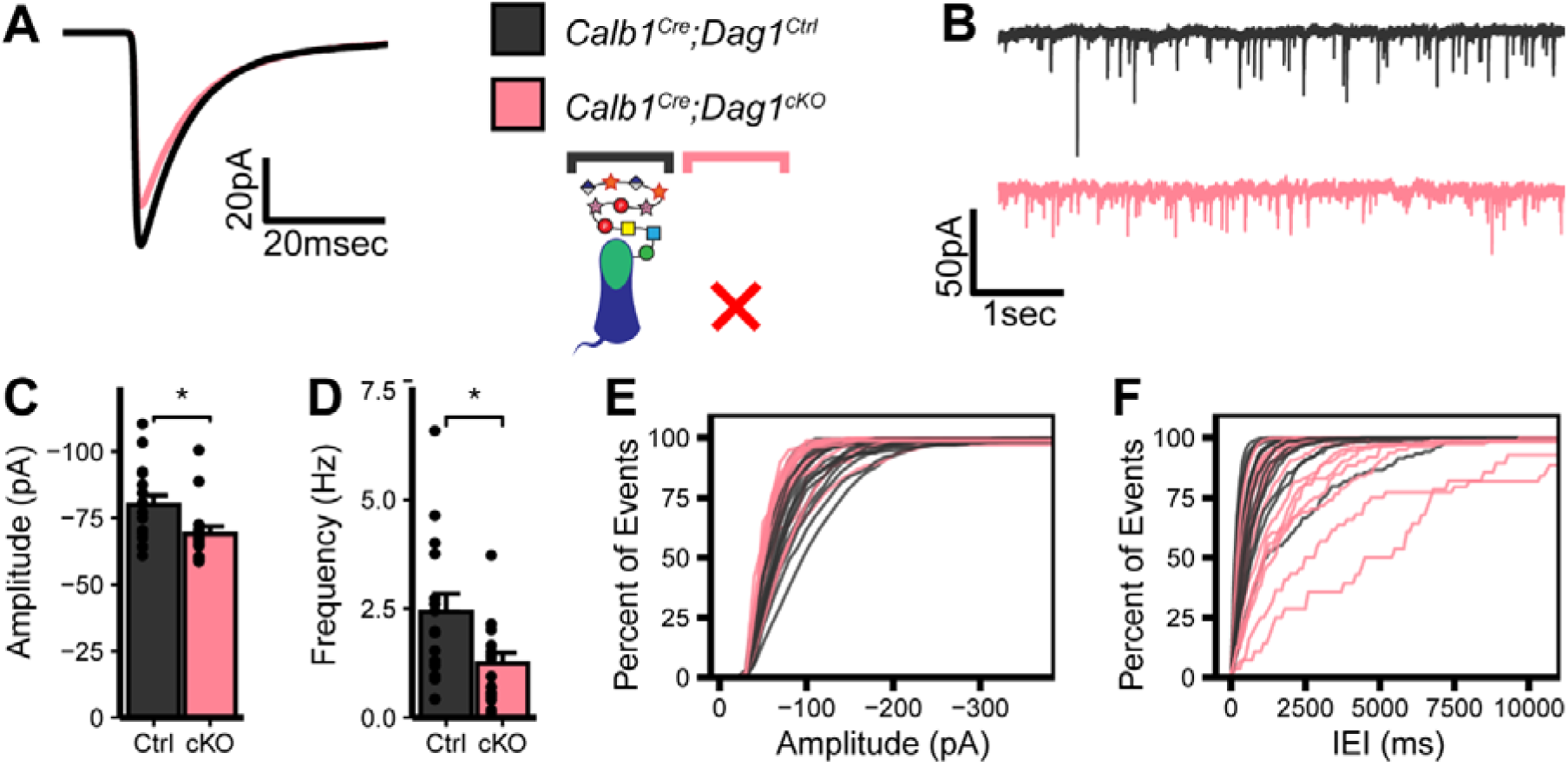
Impaired inhibitory synapse function in *Calb1^Cre^;Dag1^cKO^* Purkinje cells. **(A)** Average mIPSC event. **(B)** Representative 15 second trace of mIPSCs. **(C-D)** mIPSC amplitude **(C)** and frequency **(D)**; error bars are presented as mean + SEM. **(E-F)** Cumulative frequency histogram for individual mIPSC amplitudes **(E)** and inter-event intervals (IEI) **(F)**. *Calb1^Cre^;Dag1^Ctrl^*N = 16 cells, 6 animals. *Calb1^Cre^;Dag1^cKO^* N = 16 cells, 6 animals.

To determine whether the synaptic deficits could be explained by fewer MLIs, we immunostained lobules V/VI of cerebellar vermis for Calbindin (Purkinje cells) and Parvalbumin (Purkinje cells and MLIs) and quantified Purkinje and MLI cell densities.

**Table 1.** mIPSC rise and decay kinetics. Average mIPSC rise and decay times in milliseconds. Expressed as mean ± SEM. Ns are indicated in corresponding figure legends.

| Experiment | Rise<sub>Ctrl</sub><br>(ms) | Rise<sub>CKO</sub><br>(ms) | Rise<sub>p-val</sub> | Decay<sub>Ctrl</sub><br>(ms) | Decay<sub>CKO</sub><br>(ms) | Decay<sub>p-val</sub> |
| --- | --- | --- | --- | --- | --- | --- |
| *Calb1<sup>Cre</sup>;Dag1* | 0.73±0.05 | 0.79±0.05 | 0.4 | 7.87±0.39 | 8.77±0.62 | 0.23 |
| *Pcp2<sup>Cre</sup>;Dag1* (P25) | 1.03±0.12 | 0.97±0.11 | 0.72 | 8.64±1.32 | 8.65±0.96 | 0.99 |
| *Pcp2<sup>Cre</sup>;Dag1* (P60) | 0.93±0.1 | 0.96±0.07 | 0.82 | 7.59±0.55 | 7.78±0.55 | 0.81 |
| *Calb1<sup>Cre</sup>;Pomt2* | 0.71±0.05 | 0.72±0.05 | 0.89 | 7.61±0.58 | 7.47±0.53 | 0.86 |
| *Calb1<sup>Cre</sup>;Dag1<sup>ΔICD</sup>* | 0.66±0.06 | 0.87±0.16 | 0.23 | 7.12±0.43 | 7.55±0.84 | 0.65 |

There was no significant difference in Purkinje cell density, MLI density, or the ratio of MLIs to Purkinje cells (**Supplemental Figure 2 A-B**), suggesting that the changes in synapse function were not due to defects in cellular proliferation or survival.

To evaluate potential structural correlates of the observed reduction in mIPSC frequency and amplitude, we immunostained for the presynaptic inhibitory synapse marker VGAT along with the inhibitory postsynaptic GABA_A_ receptor subunit GABA_A_α1. Analysis of synaptic puncta in *Calb1^Cre^;Dag1^cKO^* mice using confocal microscopy with linear Wiener filter deconvolution revealed a reduction in both VGAT and GABA_A_α1 puncta density compared to littermate controls (**Figure 3 A-C**), supporting a reduction in synapse number in the absence of Dystroglycan. We also quantified synaptic puncta size and observed no change in VGAT puncta size but a significant increase in GABA_A_α1 puncta size (**Figure 3 D-E**), suggesting that Dystroglycan regulates the organization of the inhibitory postsynapse. The increase in puncta size may reflect a deficit in GABA_A_α1 receptor clustering, with GABA_A_ receptors becoming less tightly localized to the appropriate subsynaptic domains. This disruption in the alignment of synaptic microdomains could explain the observed reduction in mIPSC amplitude (Olah et al., 2023), however it does not rule out the contribution of pre-synaptic changes (Wu et al., 2022).

**Figure 3.**
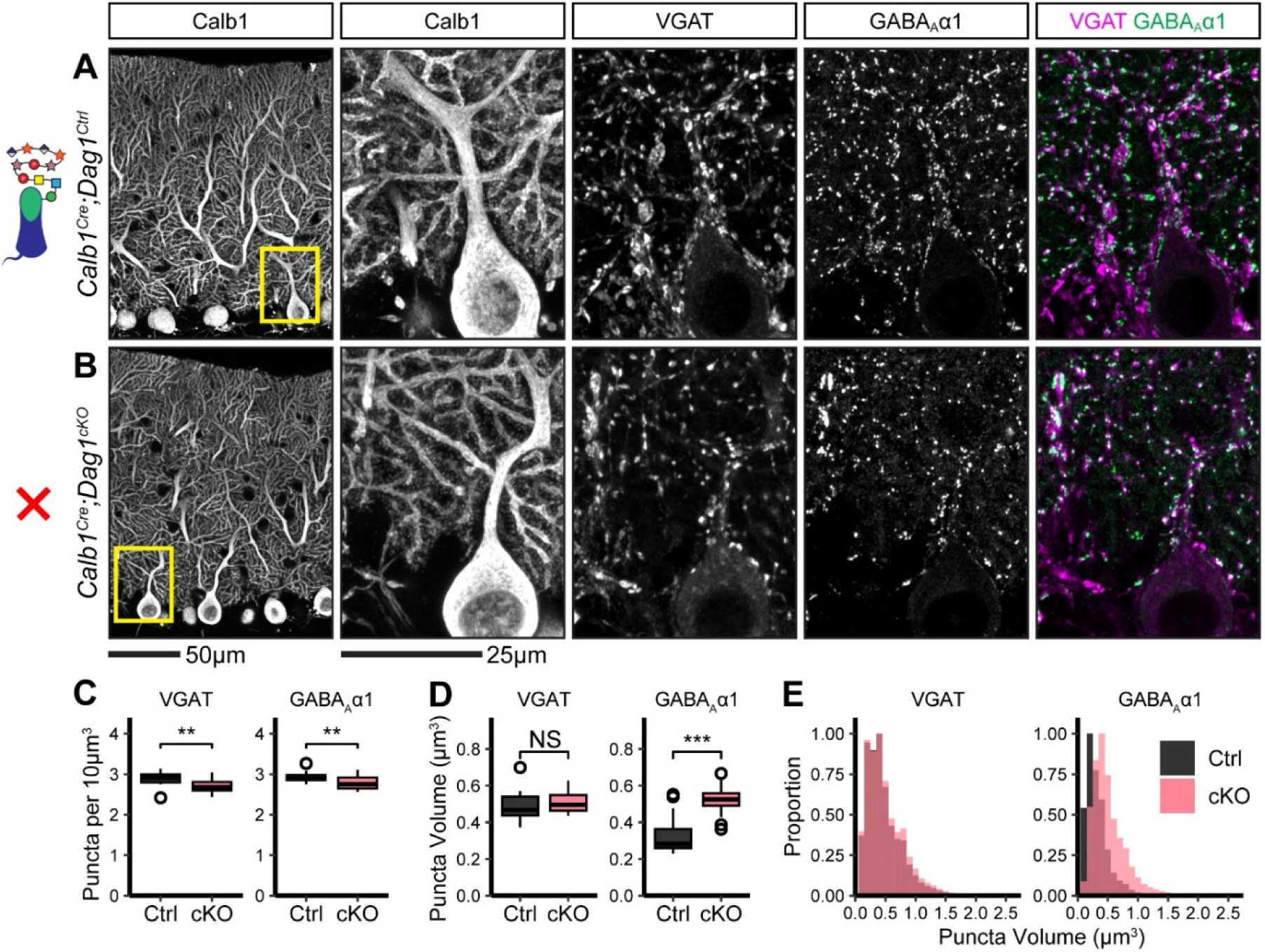
Inhibitory pre- and post-synaptic markers are altered in *Calb1^Cre^;Dag1^cKO^* cerebellar cortex. **(A-B)** Tissue sections from cerebellar vermis of *Calb1^Cre^;Dag1^cKOs^***(B)** and littermate controls **(A)** was immunostained for Calbindin to visualize Purkinje cells along with the presynaptic marker VGAT (magenta) and postsynaptic GABA_A_ receptor subunit GABA_A_α1 (green). The first panel shows a low-magnification view the Calbindin channel. Subsequent panels are magnified views of the yellow outlined region in the first panel. Images are maximum projections. **(C)** Quantification of VGAT (left) and GABA_A_α1 (right) puncta density (per 10μm^3^). Open points indicate statistical outliers, however no datapoints were excluded from statistical analysis. **(D)** Quantification of VGAT (left) and GABA_A_α1 (right) puncta size (in μm^3^). **(E)** Quantification of the distribution of VGAT (left) and GABA_A_α1 (right) puncta sizes (in μm^3^), normalized to a maximum of 1. *Calb1^Cre^;Dag1^Ctrl^* N = 20 images, 5 animals. *Calb1^Cre^;Dag1^cKO^* N = 20 images, 5 animals.

### Dystroglycan is required for the long term maintenance of inhibitory MLI:PC synapses

The synaptic phenotypes observed following early deletion of *Dag1* from Purkinje cells using *Calb1^Cre^* suggests that Dystroglycan is required for initial MLI axonal recognition of postsynaptic sites and subsequent synapse formation, similar to what we have reported for CCK^+^/CB_1_R^+^ basket synapses in the hippocampus. Testing whether Dystroglycan is involved in the maintenance of synapses requires deleting *Dag1* after synapse formation. MLI:PC synapses begin forming around P7 and are largely formed by P14, with additional synaptogenesis extending past P21 (Morales and Hatten, 2006; Viltono et al., 2008; Wizeman et al., 2019). Recent work using *L7^Cre^* to delete *Dag1* found that Dystroglycan was gradually eliminated from Purkinje cells and wasn’t fully absent from all Purkinje cells until P90. This prolonged localization of Dystroglycan at synapses in these mice likely reflects its slow rate of protein turnover as Dystroglycan has a half-life of ∼25 days in skeletal muscle (Novak et al., 2021). *L7^Cre^;Dag1^cKOs^* did exhibit impaired MLI:PC synapse function and a reduction in pre- and post-synaptic markers at P180, suggesting that Dystroglycan does indeed play a role in synapse maintenance (Briatore et al., 2020). To confirm this result, we used *Pcp2^Cre^*mice, in which Purkinje cells begin to express *Cre* gradually between P7-P14, to delete *Dag1* after synapse formation has initiated (Jahncke and Wright, 2024). To reduce the amount of Dystroglycan protein that has to be turned over, we generated compound mutants with one copy of *Dag1* constitutively deleted and the other copy flanked by *LoxP* sites for conditional deletion by *Cre*. *Pcp2^Cre^;Dag1^flox/-^*conditional knockouts (*Pcp2^Cre^;Dag1^cKO^*) showed *Cre*-mediated recombination in all Purkinje cells at P14 and loss of Dystroglycan protein by P30 (Jahncke and Wright, 2024).

The observed *Pcp2^Cre^* recombination timeline is closely aligned with the synaptogenic period in Purkinje cells and, given the timecourse of protein turnover, is likely deleting *Dag1* soon after synapse formation. When we recorded mIPSCs from *Pcp2^Cre^;Dag1^cKOs^* aged P25, they showed no significant difference from littermate controls (**Figure 4 A-F**), suggesting that synapses remain functional without Dystroglycan in the short term. However, when we recorded mIPSCs from animals aged P60, we observed a reduction in mIPSC amplitude and frequency (**Figure 4 G-L**). Together with the data from *Calb1^Cre^;Dag1^cKOs^* (**Figure 2, 3**) these results show that that Dystroglycan is required for both the formation and the long term maintenance of MLI:PC synapses.

**Figure 4.**
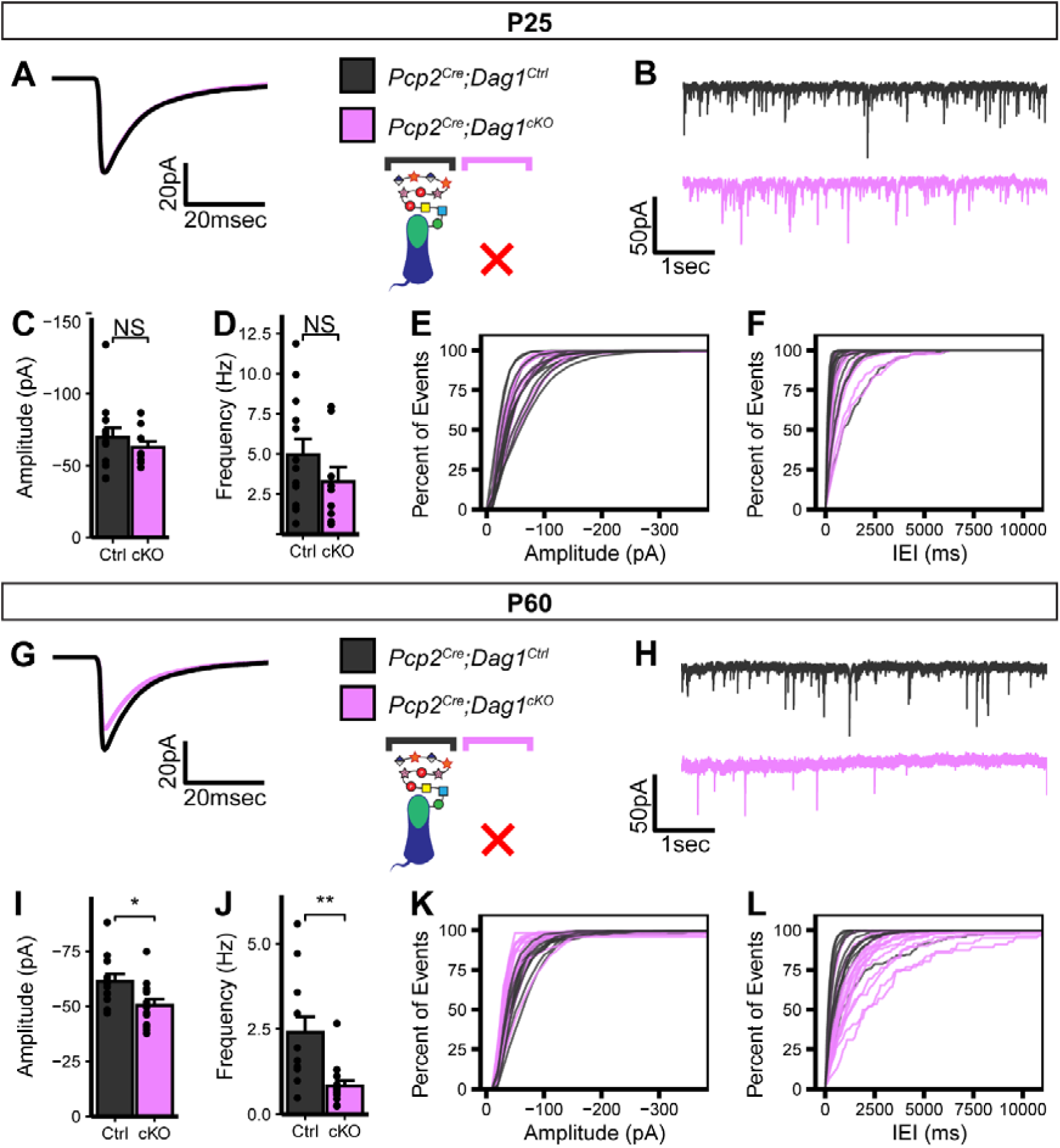
Impaired inhibitory synapse function in *Pcp2^Cre^;Dag1^cKO^* Purkinje cells at P60 but not P25. **(A-F)** mIPSC analysis in *Pcp2^Cre^;Dag1^cKO^* and littermate control Purkinje cells at P25. **(A)** Average mIPSC event. **(B)** Representative 15 second trace of mIPSCs. **(C-D)** mIPSC amplitude **(C)** and frequency **(D)**; error bars are presented as mean + SEM. **(E-F)** Cumulative frequency histogram for individual mIPSC amplitudes **(E)** and inter-event intervals (IEI) **(F)**. **(G-L)** mIPSC analysis in *Pcp2^Cre^;Dag1^cKO^* and littermate control Purkinje cells at P60. **(G)** Average mIPSC event. **(H)** Representative 15 second trace of mIPSCs. **(I-J)** mIPSC amplitude **(I)** and frequency **(J)**; error bars are presented as mean + SEM. **(K-L)** Cumulative frequency histogram for individual mIPSC amplitudes **(K)** and inter-event intervals (IEI) **(L)**. P25 *Pcp2^Cre^;Dag1^Ctrl^* N = 13 cells, 5 animals. P25 *Pcp2^Cre^;Dag1^cKO^* N = 9 cells, 5 animals. P60 *Pcp2^Cre^;Dag1^Ctrl^* N = 12 cells, 5 animals. P60 *Pcp2^Cre^;Dag1^cKO^* N = 14 cells, 4 animals.

### Extracellular glycosylation of **α**-Dystroglycan is required for inhibitory synapse formation and function

The observed synaptic phenotypes in the absence of Dystroglycan are likely driven by a combination of (1) extracellular glycan-protein interactions mediated by the matriglycan chains on α-Dystroglycan interacting with binding partners in *cis* or *trans* and (2) intracellular protein-protein interactions or signaling cascades through the intracellular C-terminus of β-Dystroglycan. To define which aspects of Dystroglycan’s synaptic function is mediated by these distinct molecular mechanisms, we generated mice with Dystroglycan either (1) lacking extracellular glycosylation or (2) lacking the intracellular domain.

The extracellular alpha subunit of Dystroglycan contains *O*-mannosyl linked glycan chains with terminal -3Xylα1-3GlcAβ1-disaccharide repeats termed “matriglycan”, which interacts with LG domain-containing extracellular binding partners (Goddeeris et al., 2013; Yoshida-Moriguchi and Campbell, 2015). The glycosyltransferase Pomt2 (Protein O-mannosyltranferase 2), in a heterocomplex with Pomt1, is responsible for adding the initial *O*-mannose to Dystroglycan, and its deletion results in a complete loss of matriglycan chains (Manya et al., 2004). We used *Calb1^Cre^* to conditionally delete *Pomt2* from Purkinje cells, generating *Calb1^Cre^;Pomt2^flox/flox^*conditional knockouts (*Calb1^Cre^;Pomt2^cKO^*) and *Calb1^Cre^;Pomt2^flox/+^*littermate controls (*Calb1^Cre^;Pomt2^Ctrl^*). Loss of Dag1 glycosylation was confirmed by immunostaining with the IIH6 antibody, which recognizes matriglycan. In contrast to the punctate synaptic staining pattern seen in *Calb1^Cre^;Pomt2^Ctrl^*Purkinje cells, *Calb1^Cre^;Pomt2^cKO^* Purkinje cells did not show any synaptic IIH6 immunoreactivity (**Supplemental Figure 3A).** We tested several potential antibodies against Dystroglycan core protein epitopes and β-Dystroglycan to confirm that loss of the glycan chains did not affect Dystroglycan’s synaptic localization, but we were unable to find immunostaining conditions in which there was specific labeling in control tissue.

To assess MLI:PC synapse function in the absence of matriglycan, we conducted mIPSC recordings in P25 *Calb1^Cre^;Pomt2^cKOs^* Purkinje cells and littermate controls (**Figure 5 A-F**). Similar to *Dag1* conditional knockouts, *Calb1^Cre^;Pomt2^cKOs^* exhibited a reduced mIPSC frequency compared to controls (**Figure 5 A-B, D, F**). However, mIPSC amplitude was normal (**Figure 5 C, E**). Based on the reduced mIPSC frequency, we hypothesized that we would observe a reduction in inhibitory synapse density in *Calb1^Cre^;Pomt2^cKOs^*similar to that observed in *Calb1^Cre^;Dag1^cKOs^*. Indeed, both VGAT and GABA_A_α1 puncta densities were reduced in *Calb1^Cre^;Pomt2^cKOs^*compared to littermate controls (**Figure 6 A-C**). We also observed an increase in GABA_A_α1, but not VGAT, puncta size, similar to that observed in the *Dag1* conditional knockouts (**Figure 6 D-E**). As the *Pomt2* conditional knockouts did not exhibit a change in mIPSC amplitude (**Figure 5**), the increased GABA_A_α1 puncta size does not appear to be driving the amplitude reduction. However, it is clear that extracellular glycan-protein interactions are involved in regulating the size and organization of the inhibitory post-synapse.

**Figure 5.**
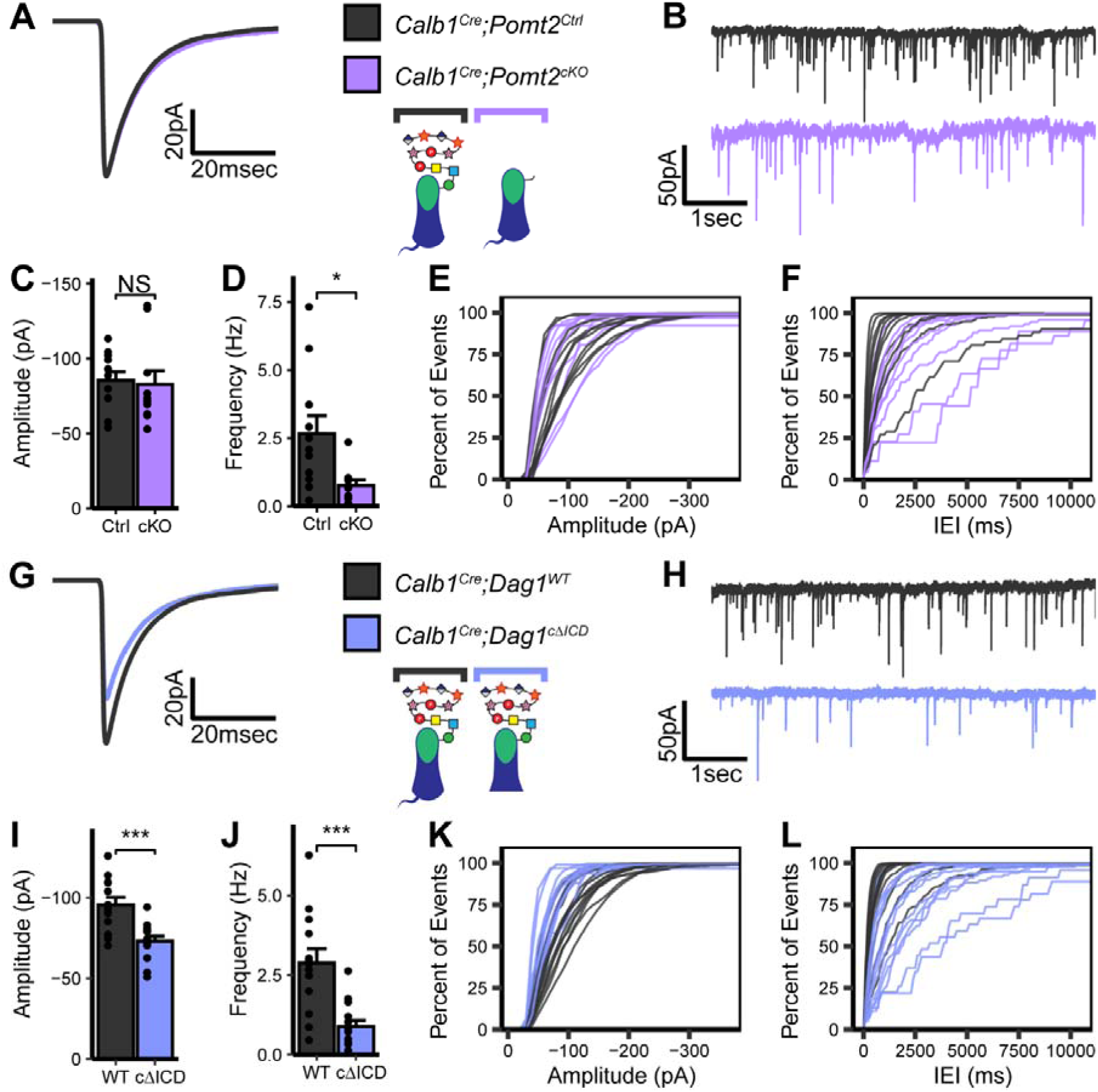
Impaired inhibitory synapse function in *Calb1^Cre^;Pomt2^cKO^* and *Calb1^Cre^;Dag1^c^*^Δ*ICD*^ Purkinje cells. **(A-F)** mIPSC analysis in *Calb1^Cre^;Pomt2^cKO^* and littermate control Purkinje cells at P25. **(A)** Average mIPSC event. **(B)** Representative 15 second trace of mIPSCs. **(C-D)** mIPSC amplitude **(C)** and frequency **(D)**; error bars are presented as mean + SEM. **(E-F)** Cumulative frequency histogram for individual mIPSC amplitudes **(E)** and inter-event intervals (IEI) **(F)**. **(G-L)** mIPSC analysis in *Calb1^Cre^;Dag1^c^*^Δ*ICD*^ and littermate control Purkinje cells at P25. **(G)** Average mIPSC event. **(H)** Representative 15 second trace of mIPSCs. **(I-J)** mIPSC amplitude **(I)** and frequency **(J)**; error bars are presented as mean + SEM. **(K-L)** Cumulative frequency histogram for individual mIPSC amplitudes **(K)** and inter-event intervals (IEI) **(L)**. *Calb1^Cre^;Pomt2^Ctrl^* N = 11 cells, 3 animals. *Calb1^Cre^;Pomt2^cKO^* N = 10 cells, 4 animals. *Calb1^Cre^;Dag1^WT^* N = 13 cells, 4 animals. *Calb1^Cre^;Dag1^c^*^Δ*ICD*^ N = 14 cells, 4 animals.

**Figure 6.**
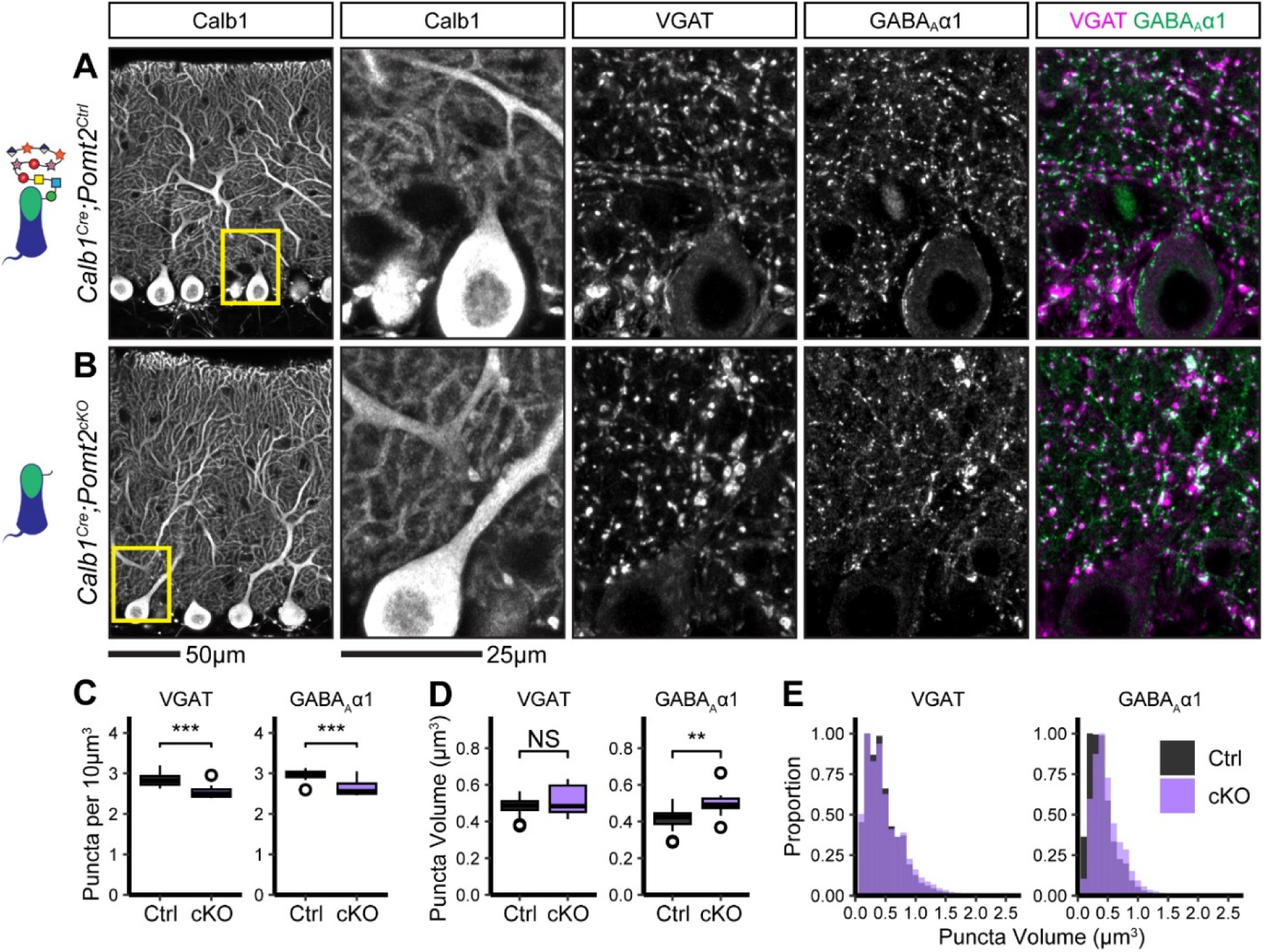
Inhibitory pre- and post-synaptic markers are altered in *Calb1^Cre^;Pomt2^cKO^* cerebellar cortex. **(A-B)** Tissue sections from cerebellar vermis of *Calb1^Cre^;Pomt2^cKOs^* **(B)** and littermate controls **(A)** was immunostained for Calbindin to visualize Purkinje cells along with the presynaptic marker VGAT (magenta) and postsynaptic GABA_A_ receptor subunit GABA_A_α1 (green). The first panel shows a low-magnification view of the Calbindin channel. Subsequent panels are magnified views of the yellow outlined region in the first panel. Images are maximum projections. **(C)** Quantification of VGAT (left) and GABA_A_α1 (right) puncta density (per 10μm^3^). Open points indicate statistical outliers, however no datapoints were excluded from statistical analysis. **(D)** Quantification of VGAT (left) and GABA_A_α1 (right) puncta size (in μm^3^). **(E)** Quantification of the distribution of VGAT (left) and GABA_A_α1 (right) puncta sizes (in μm^3^), normalized to a maximum of 1. *Calb1^Cre^;Pomt2^Ctrl^* N = 16 images, 4 animals. *Calb1^Cre^;Pomt2^cKO^* N = 12 images, 3 animals.

### The intracellular domain of **β**-Dystroglycan is required for inhibitory synapse function but not formation

The intracellular domain of Dystroglycan interacts directly with Dystrophin, which mediates interactions with the actin cytoskeleton and other scaffolding/signaling molecules. In a mouse model lacking the relevant brain isoforms of Dystrophin (*mdx*), Purkinje cells show reduced mIPSC frequency and amplitude, reduced GABA_A_α1 receptor puncta density, and fewer MLI:PC contacts, showing that Dystrophin is required for inhibitory Purkinje cell synapses in a manner similar to Dystroglycan (Anderson et al., 2003; Grady et al., 2006; Knuesel et al., 1999; Kueh et al., 2011, 2008; Wu et al., 2022). Mice lacking the cytoplasmic portion of Dystroglycan that interacts with Dystrophin develop muscular dystrophy but exhibit largely normal brain development (Satz et al., 2010). However, cytoplasmic deletion mutants exhibit synaptic deficits at CCK^+^/CB_1_R^+^ basket synapses in the hippocampus (Jahncke et al., 2024). We therefore hypothesized that in addition to extracellular Dystroglycan glycosylation, the intracellular domain of Dystroglycan may also be required for MLI:PC synapse function. To investigate the role of the intracellular domain of Dystroglycan specifically at MLI:PC synapses, we generated conditional cytoplasmic domain mutants by crossing *Calb1^Cre^;Dag1^flox/+^* mice with mice expressing one copy of truncated Dystroglycan lacking the intracellular domain (*Dag1*^Δ*ICD/+*^), generating *Calb1^Cre^;Dag1^flox/^*^Δ*ICD*^ mutants (*Calb1^Cre^;Dag1^c^*^Δ*ICD*^) and *Calb1^Cre^;Dag1^+/+^* littermate controls (*Calb1^Cre^;Dag1^WT^*). Immunohistochemistry with IIH6 showed that *Calb1^Cre^;Dag1^c^*^Δ*ICD*^ mutants maintained synaptic localization of glycosylated Purkinje cell Dystroglycan in the absence of the intracellular domain (**Supplemental Figure 3 B**).

To measure the contribution of the intracellular domain of Dystroglycan to MLI:PC synapse function we conducted mIPSC recordings from P25 *Calb1^Cre^;Dag1^c^*^Δ*ICD*^ Purkinje cells and those of littermate controls and observed a phenotype that resembled that of *Calb1^Cre^;Dag1^cKO^*Purkinje cell mIPSCs: a reduction in both mIPSC amplitude and frequency (**Figure 5 G-L**). We therefore conclude that the amplitude component of Dystroglycan-containing MLI:PC synapses is governed by intracellular interactions between Dystroglycan and other postsynaptic proteins (likely including Dystrophin) and/or intracellular signaling initiated by the C-terminus of β-Dystroglycan. The frequency component of this synaptic circuit, however, appears to be influenced by factors dictated by both extracellular glycan-protein interactions and intracellular protein-protein interactions or signaling pathways.

Due to the reduction in mIPSC frequency observed in *Calb1^Cre^;Dag1^c^*^Δ*ICD*^ Purkinje cells (**Figure 5 J, L**), we hypothesized that we would see a similar reduction in inhibitory synapse density in *Calb1^Cre^;Dag1^c^*^Δ*ICD*^ as that observed in *Calb1^Cre^;Dag1^cKO^* (**Figure 3 C**) and *Calb1^Cre^;Pomt2^cKO^* (**Figure 6 C**) Purkinje cells. To our surprise, we instead found no change in the density of presynaptic VGAT puncta, a slight increase in the density of postsynaptic GABA_A_α1 puncta, and no change in the size of synaptic puncta (**Figure 7 A-E**). This suggests that loss of the intracellular domain of Dystroglycan does not affect the ability of presynaptic MLIs to recognize postsynaptic Purkinje cells and form synaptic contacts, but it is required for proper postsynaptic composition and function.

**Figure 7.**
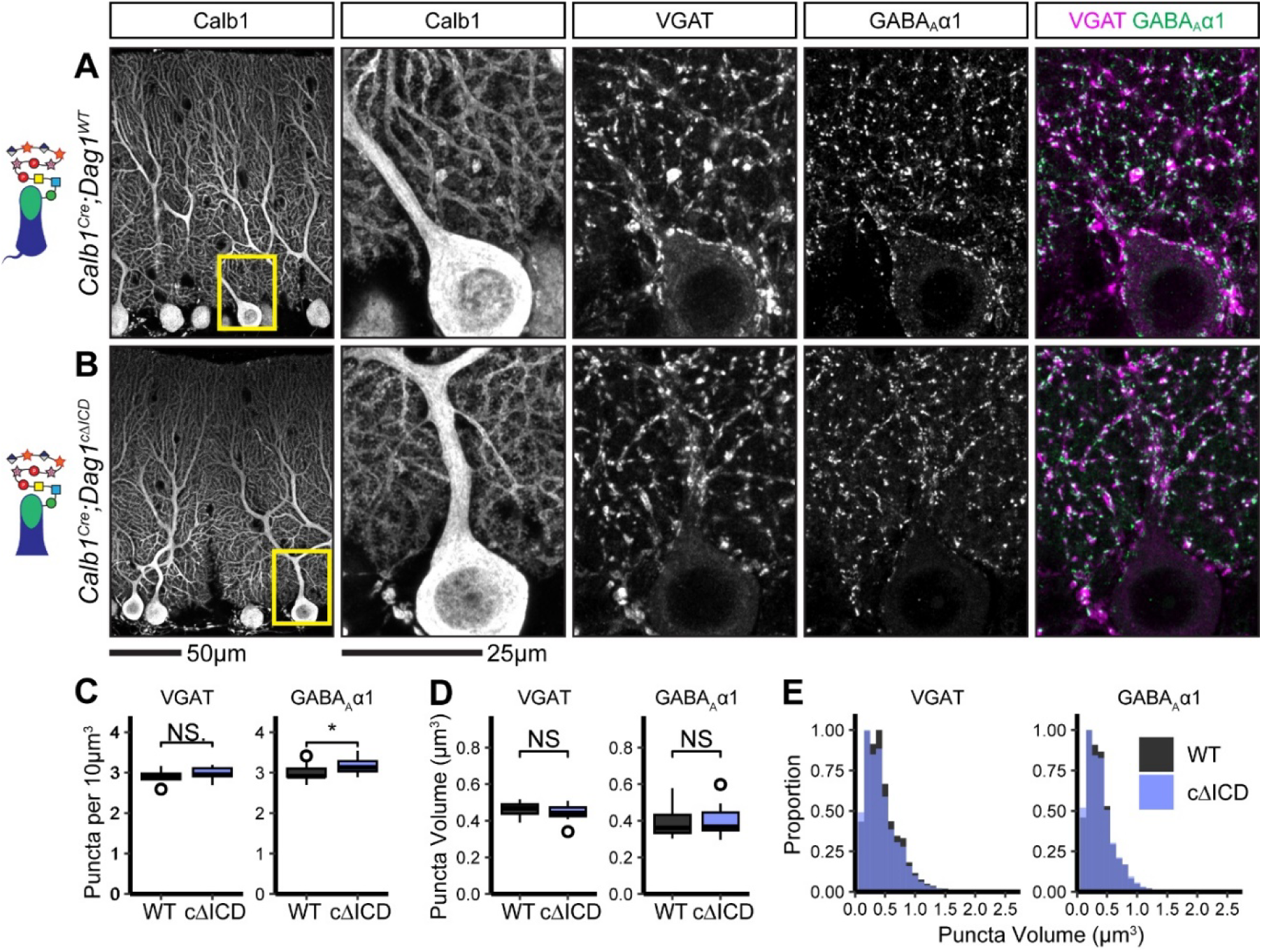
Inhibitory pre- and post-synaptic markers are relatively normal in *Calb1^Cre^;Dag1^c^*^Δ*ICD*^ cerebellar cortex. **(A-B)** Tissue sections from cerebellar vermis of *Calb1^Cre^;Dag1^c^*^Δ*ICDs*^ **(B)** and littermate controls **(A)** was immunostained for Calbindin to visualize Purkinje cells along with the presynaptic marker VGAT (magenta) and postsynaptic GABA_A_ receptor subunit GABA_A_α1 (green). The first panel shows a low-magnification view of the Calbindin channel. Subsequent panels are magnified views of the yellow outlined region in the first panel. Images are maximum projections. **(C)** Quantification of VGAT (left) and GABA_A_α1 (right) puncta density (per 10μm^3^). Open points indicate statistical outliers, however no datapoints were excluded from statistical analysis. **(D)** Quantification of VGAT (left) and GABA_A_α1 (right) puncta size (in μm^3^). **(E)** Quantification of the distribution of VGAT (left) and GABA_A_α1 (right) puncta sizes (in μm^3^), normalized to a maximum of 1. *Calb1^Cre^;Dag1^WT^* N = 13 images, 3 animals. *Calb1^Cre^;Dag1^cKO^* N = 13 images, 3 animals.

In the absence of a reduction in synapse density we are left to assume that the reduced mIPSC frequency we observed in *Calb1^Cre^;Dag1^c^*^Δ*ICD*^ Purkinje cells is due to alterations to the synapse other than simply synapse number. Previous work involving paired recordings between MLI and Purkinje cell pairs in Dystrophin-deficient *mdx* mice identified a reduction in the size of the readily releasable pool (Wu et al., 2022). A similar mechanism could explain the reduced mIPSC frequency in *Calb1^Cre^;Dag1^c^*^Δ*ICD*^ Purkinje cells (**Figure 5 J, L**). The intracellular domain of Dystroglycan interacts directly with Dystrophin, so a common mechanism between the *mdx* and *Calb1^Cre^;Dag1^c^*^Δ*ICD*^ models is likely (Jung et al., 1995). While it is clear that additional work will be necessary to determine the details underlying the observed phenotypes, these data show for the first time that the intracellular domain of Dystroglycan plays a crucial role in MLI:PC synapse function but not formation.

## Discussion

Despite years of study, the molecules required for MLI:PC synaptogenesis remain enigmatic. Several postsynaptic transmembrane proteins including Neuroligins, Calsyntenin-3, Kit, and Dystroglycan are required for proper MLI:PC synapse maintenance and function (Briatore et al., 2020; Liu et al., 2022; Pettem et al., 2013; Zaman et al., 2024; Zhang et al., 2015). However, these prior studies were unable to address the role of these proteins in initial synaptogenesis due to the lack of genetic tools that target Purkinje neurons early in development. With the identification of *Calb1^Cre^* as a tool for the deletion of genes from Purkinje cells early in development (Daigle et al., 2018; Jahncke and Wright, 2024), we were able to show for the first time that Dystroglycan is required for initial synapse formation at MLI:PC synapses (**Figures 2, 3**). Loss of Dystroglycan was accompanied by reductions in both mIPSC amplitude, mIPSC frequency, and inhibitory pre- and postsynaptic marker density, illustrating that synapses are not able to form normally in the absence of Dystroglycan.

Previous work used the *L7^Cre^* line to delete *Dag1* from Purkinje cells after synaptogenesis, which did not achieve complete loss of Dystroglycan protein until P90, limiting the study to the role of Dystroglycan in long-term synapse maintenance (Briatore et al., 2020; Jahncke and Wright, 2024). Nevertheless, it was determined that Dystroglycan is indeed required for synapse maintenance, as the *L7^Cre^;Dag1^flox/flox^* Purkinje cells showed a decreased inhibitory synapse density and decreased mIPSC frequency and amplitude (Briatore et al., 2020). We confirmed these results using a *Pcp2^Cre^*mouse line mated with mice in which only one copy of *Dag1* needed to be excised (*Dag1^flox/-^*), achieving complete loss of Dystroglycan by P30 (Jahncke and Wright, 2024). We observed similarly impaired synapse function in *Pcp2^Cre^;Dag1^flox/-^* (*Pcp2^Cre^;Dag1^cKO^*) Purkinje cells at P60 (**Figure 4 G-L**), confirming Dystroglycan’s role in MLI:PC synapse maintenance. Curiously, mIPSC recordings in *Pcp2^Cre^;Dag1^cKO^*Purkinje cells at P25, in which *Dag1* is deleted from most cells and most protein turned over, showed no significant impact on MLI:PC function (**Figure 4 A-F**). There are two potential explanations for this result: (1) synapses remain stable for some period of time (most likely on the order of days to weeks) after Dystroglycan protein is eliminated, or (2) the amount of Dystroglycan protein remaining at P25 is below the detection threshold for immunohistochemistry, but sufficient to maintain synapse function.

### Refining the molecular mechanism of Dystroglycan’s contributions to synapse function

How does Dystroglycan regulate MLI:PC synapse formation and function? The genetic manipulations we used in this study allowed us to distinguish the extracellular (*Calb1^Cre^;Pomt2^cKO^*) and intracellular (*Calb1^Cre^;Dag1^c^*^Δ*ICD*^) functions of Dystroglycan at these synapses. Our experiments using *Calb1^Cre^;Pomt2^cKO^*mice show that extracellular glycosylation of α-Dystroglycan is required for Dystroglycan’s role in both synaptogenesis and synapse function (**Figures 5, 6**), in agreement with our previous work showing that forebrain deletion of *Dag1* or *Pomt2* results in identical deficits in hippocampal CCK^+^/CB_1_R^+^ basket synapse formation and function (Jahncke et al., 2024). Removal of *Pomt2* results in loss of all *O*-mannosyl linked glycosylation on Dystroglycan (**Supplemental Figure 3**) (Manya et al., 2004), including the -3Xylα1-3GlcAβ1- disaccharide repeats referred to as matriglycan elongated by the glycosyltransferase *LARGE1* (Briggs et al., 2016; Inamori et al., 2012). The length of the matriglycan repeat can serve as a “tunable matrix” capable of binding multiple LG domain-containing proteins simultaneously, allowing large macromolecular complexes to form (Goddeeris et al., 2013; Sheikh et al., 2022). Purkinje cell matriglycan migrates at a significantly higher molecular weight than matriglycan in other neuronal and non-neuronal populations, suggesting Purkinje cells may contain longer matriglycan chains with increased binding capacity (Satz et al., 2010).

The intracellular domain of Dystroglycan was long thought to be dispensable for neuronal function, as mice lacking the intracellular domain of Dystroglycan exhibit normal neuronal migration and axon targeting (Jahncke et al., 2024; Lindenmaier et al., 2019; Satz et al., 2010). However, we recently found that hippocampal CCK^+^/CB_1_R^+^ basket cells exhibit slightly altered perisomatic axon targeting and impaired synapse function despite normal synapse formation when the intracellular domain is deleted (Jahncke et al., 2024). In the present study we similarly found impaired synapse function in *Calb1^Cre^;Dag1^c^*^Δ*ICD*^ mice (**Figure 5 G-L**) despite relatively normal MLI:PC synapse formation (**Figure 7**). Not only do these new findings confirm the role of the cytoplasmic domain in synapse function, but they also illustrate the cell autonomous nature of the mechanism as the *Calb1^Cre^*mutation is restricted to Purkinje cells within the cerebellum. Taken together, the results obtained in *Calb1^Cre^;Pomt2^cKO^* and *Calb1^Cre^;Dag1^c^*^Δ*ICD*^ mutants show that both extracellular glycosylation and intracellular scaffolding/signaling by Dystroglycan is required for proper MLI:PC synapse function, while the intracellular domain is dispensable for the initial formation of presynaptic contacts onto Purkinje cells.

### Dag1 extracellular interactions shape MLI:PC synapse development and function

The identity of Dystroglycan’s presynaptic partner(s) at MLI:PC synapses remains unknown. Matriglycan binds specifically to proteins with LG domains, and there are upwards of 20 LG domain-containing proteins expressed in cerebellar cellular populations that are candidate extracellular binding partners of Dystroglycan. Biochemical isolation of Dystroglycan-containing complexes to identify potential synaptic binding partners is complicated by its expression in many cell types in the brain, with neuronal Dystroglycan representing a minority proportion. Possible binding partners for Dystroglycan at MLI:PC synapses include members of the Neurexin family of presynaptic adhesion molecules. Biochemical studies show that Dystroglycan binds to two of the LG domains present in Neurexins in a glycosylation-dependent manner, and Dystroglycan:Neurexin-3 interactions regulate inhibitory synapse function in the olfactory bulb and medial prefrontal cortex (Reissner et al., 2014; Sugita et al., 2001; Trotter et al., 2023). However, whether this interaction occurs at MLI:PC synapses has not yet been examined. All three Neurexins are expressed by MLIs, and these neurons have been difficult to selectively target for conditional deletions (Kozareva et al., 2021; Saunders et al., 2018).

Neurexins also interact with postsynaptic Neuroligins, and all three Neuroligin isoforms are expressed in Purkinje cells. Triple conditional deletion of *Neuroligin1/2/3* from Purkinje cells causes an increase in inhibitory MLI:PC synaptic puncta size and reduced mIPSC frequency and amplitude, similar to our observations when *Dystroglycan* is conditionally deleted from Purkinje cells (**Figure 2, 3**) (Zhang et al., 2015). This effect was not observed in conditional *Nlgn1/Nlgn3* double knockouts but was seen in *Ngln2/Nlgn3* double knockout. A *Nlgn2* constitutive knockout only partially recapitulated these results, suggesting that while Nlgn2 is the primary driver of this phenotype, some degree of molecular compensation can occur in its absence (Zhang et al., 2015). Neuroligins are thought to be linked to the intracellular domain of β-Dystroglycan through S-SCAM/MAGI-2 (Sumita et al., 2007), and Nlgn2 puncta density is reduced at somatic MLI:PC synapses in Purkinje cell *Dag1* conditional knockouts (Briatore et al., 2020; Nguyen and Südhof, 1997). Thus, Dystroglycan can interact directly and indirectly with Neurexins and Neuroligins in a network of pre- and post-synaptic cell adhesion complexes.

Synapse formation occurs in multiple phases: a presynaptic axon must first recognize its postsynaptic target through some sort of recognition cue, after which pre- and post-synaptic molecules are recruited to the nascent synapse. While presynaptic Neurexins and postsynaptic Neuroligins are undoubtedly critical for the maturation and maintenance of functional MLI:PC synapses, neither are generally thought to be required for initial synaptogenesis. Constitutive or developmentally early deletions of Neurexins or Neuroligins typically result in impaired synapse function without a decrease in synapse number, suggesting that initial postsynaptic partner recognition and synaptogenesis is mediated by other molecules (Liang et al., 2015; Missler et al., 2003; Trotter et al., 2023; Varoqueaux et al., 2006; Zhang et al., 2015). Dystroglycan, on the other hand, is required for both the initial formation of a synapse and its maintenance; developmentally early deletion of *Dag1* results in a loss of synapses and reduced synaptic function (**Figure 2, 3**) (Früh et al., 2016; Jahncke et al., 2024). Similarly, later deletion of *Dag1* also impacts synapse function, showing it is also required for subsequent synapse maintenance (**Figure 4**) (Briatore et al., 2020).

Together with evidence from other Dystroglycan-expressing synapses in the brain (Früh et al., 2016; Jahncke et al., 2024; Miller and Wright, 2021; Trotter et al., 2023), our data suggests that Dystroglycan acts as a recognition molecule for presynaptic axon targeting of the postsynaptic compartment, after which its role switches to one of postsynaptic molecule recruitment and synapse stabilization. While the glycan-protein interaction between Dystroglycan and Neurexin may be required for the maturation and stabilization of a synapse, the presynaptic binding partner(s) of Dystroglycan during axon targeting may be mediated by other LG domain-containing proteins.

### Dystroglycan’s intracellular domain is required for inhibitory synapse function

In both the hippocampus (Jahncke et al., 2024) and the cerebellum (**Figures 5, 7**), the intracellular domain of Dystroglycan is required for postsynaptic function, while presynaptic targeting is largely normal. While intracellular interactions with Dystroglycan in the brain remain poorly described, the prevailing interaction contributing to MLI:PC synapse function is likely between Dystroglycan and Dystrophin (Dmd). Dystroglycan and Dystrophin interact directly through the intracellular domain of Dystroglycan (Rosa et al., 1996; Suzuki et al., 1994). There is extensive literature supporting a role for Dystrophin at MLI:PC synapses (Anderson et al., 2003; Briatore et al., 2020; Gao and McNally, 2015; Knuesel et al., 1999; Kueh et al., 2011, 2008; Wu et al., 2022). Similar to our data in *Calb1^Cre^;Dag1^c^*^Δ*ICD*^ Purkinje cells (**Figure 5 G-L**), Dystrophin-deficient *mdx* mice exhibit impaired mIPSC frequency and amplitude (Anderson et al., 2003; Kueh et al., 2011, 2008; Wu et al., 2022). Since *mdx* mice have a constitutive mutation in Dystrophin, it is not clear if it functions in Purkinje cells, MLIs, or both. Dystrophin is predominantly localized to the postsynaptic compartment at MLI:PC synapses (Lidov et al., 1990), and its loss results in abnormal GABA receptor clustering in Purkinje cells (Knuesel et al., 1999), suggesting a postsynaptic role. However, loss of Dystrophin also affects presynaptic MLI function. Paired recordings between MLIs and PCs in *mdx* mice revealed an increase in the failure rate, but no change in paired pulse ratio, suggesting that release probability does not explain the increased failure rate (Wu et al., 2022). Rather, the size of the readily releasable pool and the quantal content were both reduced in *mdx* MLIs (Wu et al., 2022). One possibility is that Dystroglycan acts as the link between postsynaptic Dystrophin and presynaptic defects in MLI function, possibly through transsynaptic interactions with Neurexins. Studies using MLI:PC paired recordings in *Calb1^Cre^;Dag1^c^*^Δ*ICD*^ mice or Purkinje cell specific deletion of *Dystrophin* could inform whether this proposed mechanism is at play.

### Molecular diversity among MLI:PC synapses

Historically MLIs have been classified based on their morphological features, with basket cells in the lower third of the molecular layer targeting their axons to Purkinje cell somata and proximal dendrites while stellate cells located in the upper third of the molecular layer target their axons to Purkinje cell distal dendrites. However, recent studies have blurred these distinctions. While transcriptomic classification identified two distinct MLI populations, both types were present throughout the entire molecular layer and did not precisely correspond to the classical morphological distinction of basket and stellate cells (Kozareva et al., 2021; Wang and Lefebvre, 2022). Subsequent functional comparisons showed that the MLI1 transcriptional subtype primarily forms inhibitory inputs onto Purkinje cells, while the MLI2 transcriptional subtype primary forms inhibitory inputs onto MLI1 cells (Lackey et al., 2024).

IIH6 immunoreactivity shows Dystroglycan localized at synaptic puncta on Purkinje cell somata and dendrites throughout the entirety of the molecular layer (Error! Reference source not found.). Deletion of *Dystroglycan* (Jahncke and Wright, 2024) or *Pomt2* (**Supplemental Figure 3A**) from Purkinje cells with *Calb1^Cre^*results in a complete loss of punctate IIH6 staining in the molecular layer, suggesting Dystroglycan is predominantly localized at MLI1:PC synapses. While loss of Dystroglycan caused a reduction in both VGAT and GABA_A_α1 puncta throughout the Purkinje cell layer and molecular layer, the reduction on the surface of the Purkinje cell soma was particularly striking, especially so for the postsynaptic GABA_A_α1 receptor subunit (**Figure 3 A-B**). One unique molecular feature of somatic MLI:PC synapses is that they lack the inhibitory postsynaptic scaffolding molecule Gephyrin, whereas dendritic MLI:PC synapses do contain Gephyrin (Viltono et al., 2008). While GABA_A_ receptors are typically thought to be anchored to the post-synapse through Gephyrin, Dystroglycan may play a more pronounced role in scaffolding postsynaptic receptors at Gephyrin-lacking somatic MLI:PC synapses. A complete accounting of the molecular composition of Dystroglycan-containing postsynaptic inhibitory complexes, and how this may be different in distinct subcellular compartments in Purkinje cells remains an open question.

## Materials and Methods

### Animal Husbandry

All animals were housed and cared for by the Department of Comparative Medicine (DCM) at Oregon Health and Science University (OHSU), an AAALAC-accredited institution. Animal procedures were approved by OHSU Institutional Animal Care and Use Committee (Protocol # IS00000539), adhered to the NIH *Guide for the care and use of laboratory animals*, and provided with 24-hour veterinary care. Animal facilities are regulated for temperature and humidity and maintained on a 12-hour light-dark cycle and were provided food and water *ad libitum*. Mice were used between ages P21-P60 (as indicated in the text or figure legend). Mice were euthanized by administration of CO_2_ followed by exsanguination.

### Mouse Strains and Genotyping

The day of birth was designated postnatal day 0 (P0). Ages of mice used for each analysis are indicated in the figure and figure legends. Mouse strains used in this study have been previously described and were obtained from Jackson Labs, unless otherwise indicated (Error! Reference source not found.). Breeding schemas are as described in Error! Reference source not found.. Where possible, mice were maintained on a C57BL/6 background. The *Dag1*^Δ*ICD*^ line was outcrossed to a CD-1 background for one generation to increase the viability of mutant pups. The *Dag1*^Δ*ICD*^ line was then backcrossed to C57BL/6 breeders for 3 generations before performing experiments.

*Dag1^+/-^* mice were generated by crossing the *Dag1^flox/flox^*line to a *Sox2^Cre^* line to generate germline *Dag1*^Δ*/+*^ mice hereafter referred to as *Dag1^+/-^* as the resultant transcript is nonfunctional. These mice were thereafter maintained as heterozygotes. Genomic DNA extracted from toe or tail samples using the HotSHOT method (Truett et al., 2000) was used to genotype animals. Primers for genotyping can be found on the JAX webpage or originating article. *Dag1^+/-^* mice were genotyped with the following primers: CGAACACTGAGTTCATCC (forward) and CAACTGCTGCATCTCTAC (reverse). For each mouse strain, littermate controls were used for comparison with mutant mice. For all experiments mice of both sexes were used indiscriminately.

**Table 2.**
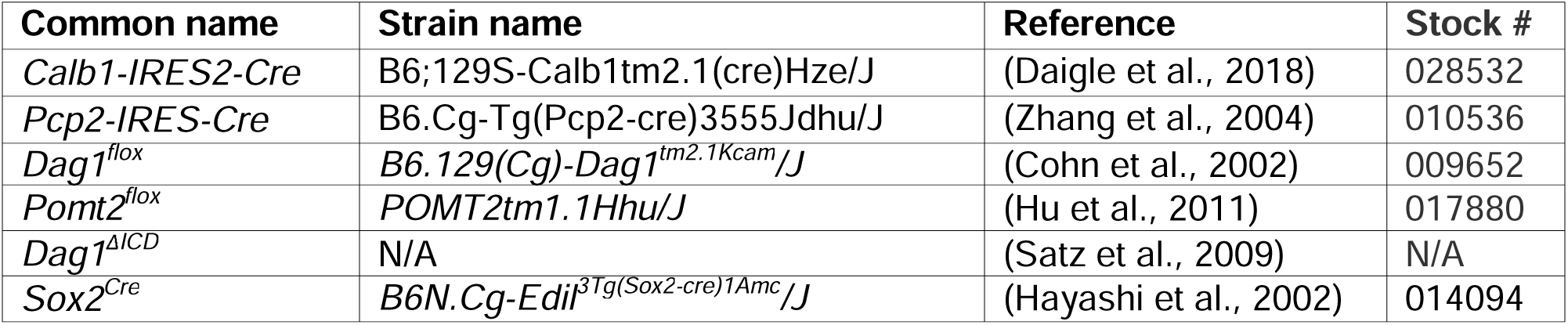
Mouse strains.

**Table 3.** Breeding schemes.

| Breeding Scheme | Control Genotype | Mutant Genotype |
| --- | --- | --- |
| *Calb1<sup>Cre/+</sup>;Dag1<sup>+/-</sup>* x *Dag1<sup>flox/flox</sup>* | *Calb1<sup>Cre/+</sup>;Dag1<sup>flox/+</sup>* | *Calb1<sup>Cre/+</sup>;Dag1<sup>flox/-</sup>* |
| *Pcp2<sup>Cre/+</sup>;Dag1<sup>+/-</sup>* x *Dag1<sup>flox/flox</sup>* | *Pcp2<sup>Cre/+</sup>;Dag1<sup>flox/+</sup>* | *Pcp2<sup>Cre/+</sup>;Dag1<sup>flox/-</sup>* |
| *Calb1<sup>Cre/+</sup>;Pomt2<sup>flox/+</sup>* x *POMT2<sup>flox/flox</sup>* | *Calb1<sup>Cre/+</sup>;Pomt2<sup>flox/+</sup>* | *Calb1<sup>Cre/+</sup>;Pomt2<sup>flox/flox</sup>* |
| *Dag1<sup>ΔICD/+</sup>* x *Calb1<sup>Cre/+</sup>;Dag1<sup>flox/+</sup>* | *Calb1<sup>Cre/+</sup>;Dag1<sup>+/+</sup>* | *Calb1<sup>Cre/+</sup>;Dag1<sup>ΔICD/flox</sup>* |

### Perfusions and tissue preparation

Mice were perfused with either 4% paraformaldehyde (PFA) in PBS, pH 7 (**Figure 1, Supplemental Figures 1, 2, 3**) or 9% Glyoxal/8% Acetic acid (GAA) in PBS, pH 4-5 (modified from Konno et al., 2023) (**Figures 3**, **6**, **7**). PFA was prepared from powder (Thermo Scientific Chemicals, Cat. No. A1131336). Glyoxal was purchased as a 40% stock solution (Thermo Scientific Chemicals, Cat. No. 156225000) and Glacial Acetic Acid was purchased from Fisher (Cat. No. A38-500). PFA perfusion was used for all immunohistochemical experiments except for those involving anti-GABA_A_α1, which was only compatible with GAA perfused tissue. P30 mice were deeply anesthetized using CO_2_ and transcardially perfused with ice cold 0.1M PBS for two minutes to clear blood from the brain, followed by either (1) 15 mL of ice cold 4% PFA in PBS or (2) 20 mL of ice cold 9%/8% GAA in PBS, as indicated. After perfusion, brains were dissected and post-fixed in either (1) 4% PFA for 30 minutes at room temperature or (2) 9%/8% GAA overnight at 4°C. Brains were rinsed with PBS, embedded in 4% low-melt agarose (Fisher, Cat. No. 16520100), and sectioned at 70μm using a vibratome (VT1200S, Leica Microsystems Inc., Buffalo Grove, IL) into 24-well plates containing 1 mL of 0.1M PBS with 0.02% Sodium Azide.

### Immunohistochemistry

Single and multiple immunofluorescence detection of antigens was performed as follows: free-floating vibratome sections (70μm) were briefly rinsed with PBS, then blocked for 1 hour in PBS containing 0.2% Triton-X (PBST) plus 2% normal goat serum. For staining of Dystroglycan synaptic puncta, an antigen retrieval step was performed prior to the blocking step: sections were incubated in sodium citrate solution (10mM Sodium Citrate, 0.05% Tween-20, pH 6.0) for 12 min at 95°C in a water bath followed by 12 min at room temperature. Sections were incubated with primary antibodies (Error! Reference source not found.) diluted in blocking solution at 4°C for 48-72 hours. Following incubation in primary antibody, sections were washed with PBS three times for 20 min each. Sections were then incubated with a cocktail of secondary antibodies (1:500, Alexa Fluor 488, 546, 647) in blocking solution containing Hoechst 33342 (1:10,000, Life Technologies, Cat. No. H3570) overnight at room temperature. Finally, sections were mounted on slides using Fluoromount-G (SouthernBiotech) and sealed using nail polish.

**Table 4.** Primary antibodies used for immunohistochemistry.

| Target | Host | Dilution | Source | Catalog # | RRID |
| --- | --- | --- | --- | --- | --- |
| α-Dystroglycan (IIH6C4) | Mouse | 1:250 | MilliporeSigma | 05-593 | AB_309828 |
| α-Dystroglycan (IIH6C4) | Mouse | 1:250 | MilliporeSigma | 05-593-I | AB_3083460 |
| VGAT | Rabbit | 1:500 | Synaptic Systems | 131-003 | AB_887869 |
| VGAT | Guinea Pig | 1:500 | Synaptic Systems | 131-005 | AB_1106810 |
| GAD67 | Mouse | 1:500 | MilliporeSigma | MAB5406 | AB_2278725 |
| VGLUT1 | Guinea Pig | 1:500 | MilliporeSigma | AB5905 | AB_2301751 |
| VGLUT2 | Guinea Pig | 1:250 | Synaptic Systems | 135-404 | AB_887884 |
| GABA<sub>A</sub> α1 | Mouse | 1:500 | NeuroMab | N95/35 | AB_2108811 |
| Parvalbumin | Rabbit | 1:1000 | Swant | PV27 | AB_2631173 |
| Parvalbumin | Goat | 1:1000 | Swant | PVG213 | AB_2650496 |
| Calbindin | Rabbit | 1:1000 | Swant | CB38 | AB_10000340 |
| Calbindin | Chicken | 1:2000 | Boster Bio | M03047-2 | AB_2936235 |

### Microscopy

Imaging was performed on either a Zeiss Axio Imager M2 fluorescence upright microscope equipped with an Apotome.2 module or a Zeiss LSM 980 laser scanning confocal build around a motorized Zeiss Axio Observer Z1 inverted microscope with a Piezo stage. As indicated below (Error! Reference source not found.), some experiments utilizing the LSM 980 confocal, a linear Wiener filter deconvolution step (Zeiss LSM Plus) was used at the end of image acquisition with 1.2X Nyquist sampling. The Axio Imager M2 uses a metal halide light source (HXP 200 C), Axiocam 506 mono camera, and 20X/0.8 NA Plan-Apochromat objectives. The LSM 980 confocal light path has two multi-alkali PMTs and two GaAsP PMTs for four track imaging. Confocal images were acquired using a 63X/1.4 NA Plan-Apochromat Oil DIC M27 objective. Z-stack images were acquired and analyzed offline in ImageJ/FIJI (Schindelin et al., 2012) or Imaris 10.0 (Oxford Instruments). Images used for quantification between genotypes were acquired using the same exposure times or laser power. Brightness and contrast were adjusted in FIJI to improve visibility of images for publication. Figures were composed in Adobe Illustrator 2024 (Adobe Systems).

**Table 5.** Image acquisition setup microscopy experiments.

| Microscope | Objective | Figure(s) |
| --- | --- | --- |
| Zeiss Axio Imager M2 | 20X/0.8 NA | Supplemental Figure 2 |
| Zeiss LSM 980 | 63X/1.4 NA | Figure 1, Supplemental Figure 1 |
| Zeiss LSM 980 (with Weiner deconvolution) | 63X/1.4 NA | Figures 3, 6, 7, Supplemental Figure 3 |

### Image Quantification

For imaging experiments, 4-8 images were acquired from 3-4 sagittal sections per animal, and at least three animals per genotype were used for analysis. Sections were chosen from cerebellar vermis and images were acquired within cerebellar cortex of lobules V-VI.

#### Synaptic Puncta Density, Size, and Colocalization

0.2μm z-stacks covering 5μm were acquired using a 63X objective on a Zeiss LSM 980 (as described above). Analysis of image stacks was performed in Imaris 10.0 (Oxford Instruments). The Surfaces function was used to reconstruct the Purkinje cells using the Parvalbumin (Error! Reference source not found., Error! Reference source not found.) or Calbindin (**Figures 3**, **6**, **7**) channel. To enrich for Purkinje cell signal, the synaptic marker channels were masked to only include information at the surface or inside of the Purkinje cell surface. The Spots function was used to determine the location of synaptic puncta in 3-dimensional space using local contrast to identify puncta. Spots with a volume less than 0.1μm^3^ were excluded. Puncta density (puncta per 10μm^3^) was calculated by dividing 10 by the Average Distance To 5 Nearest Neighbours. Spots were determined to be colocalized if they were within 1μm. As a control for random colocalization, one synaptic marker channel was mirrored and colocalization was recalculated.

#### Purkinje Cell and MLI Cell Densities

0.5μm z-stacks covering 20μm were acquired using a 20X objective on a Zeiss Axio Imager M2. Maximum projections were used for analysis in FIJI (Schindelin et al., 2012). Cells were counted using the Multi-Point Tool. The length of the region of cerebellar cortex analyzed was measured using the Freehand Line tool; this value was used to normalize the cell counts to unit length.

### Electrophysiology

For acute slice preparation, mice were deeply anesthetized in 4% isoflurane and subsequently injected with a lethal dose of 2% 2, 2, 2-Tribromoethanol in sterile water followed by transcardial perfusion with 10mL ice cold cutting solution containing the following (in mM): 100 Choline Chloride, 2.5 KCl, 1.2 NaH_2_PO_4_, 25 NaHCO_3_, 3 3-myo-inositol, 25 glucose, 5 Na Ascorbate, 2 Na Pyruvate, 7 MgSO_4_, 0.5 CaCl_2_; pH 7.3, 300-340mmol/kg. After rapid decapitation, the brain was briefly submerged in ice cold cut solution bubbled with carbogen (95% oxygen, 5% CO2) and then sectioned into 300μm sagittal sections (Leica VT1200S vibratome) in bubbled ice-cold cut solution. Slices were recovered in 37°C recording ACSF, bubbled, for 15 minutes followed by 1 hour in room temperature recording ACSF (in mM: 125 NaCl, 25 NaHCO_3_, 1.25 NaH_2_PO_4_, 3 KCl, 25 D-Glucose, 2 CaCl_2_, 1 MgCl_2_) with an osmolarity of 310-325mmol/kg and supplemented with 1.5mM Na Ascorbate, bubbled.

Purkinje cells were patched in whole cell configuration using 1.2-2MΩ borosilicate glass pipettes filled with high chloride internal solution containing the following (in mM): 125 CsCl, 2.5 MgCl_2_, 0.5 EGTA, 10 HEPES, 2 Mg-ATP, 0.3 Na-GTP, 5 QX-314; pH 7.2, 300mmol/kg. Pipettes were wrapped in parafilm to reduce capacitive currents. Cells were voltage clamped at −70mV and continuously superfused with 2-3 mL/min bubbled recording ACSF (310-325mmol/kg) containing 10μM NBQX to block excitatory transmission and 500nM TTX to block action potentials. After reaching a stable baseline, 5 minutes of mIPSCs were recorded. Signals were amplified with an AxoPatch 200B amplifier (Molecular Devices), low-pass filtered at 5 kHz, and digitized and sampled at 10 kHz with a NIDAQ analog-to-digital board (National Instruments). Data were acquired and analyzed using a custom script in Igor Pro 8 (Wavemetrics). A hyperpolarizing step of −10mV was applied before each sweep to monitor input resistance, series resistance, and measure cell capacitance. Series resistance was not compensated and was maintained below 20MΩ. Cells were excluded if series resistance changed by more than 25%.

Rise and decay kinetics were calculated on each detected event for a given cell and averaged to a single value for that cell. Rise time was defined as the amount of time between 10% and 90% of the maximum amplitude of a given event. Decay was calculated as the time constant when the decay phase of the event was fit to an exponential curve.

### Statistical analysis

Phenotypic analyses were conducted using tissue collected from at least three mice per genotype from at least two independent litters. The number of mice and replicates used for each analysis (“N”) are indicated in the text or figure legends. Power analysis was used to determine samples sizes with α = 0.05 and β = 0.80 and effect size determined using pilot data. Phenotypes were indistinguishable between male and female mice and were analyzed together. Analyses were performed blind to genotype. For comparisons between two groups, significance was determined using a two-tailed Students t-test. For comparisons between more than two groups, significance was determined using a 2-way ANOVA with Tukey HSD post-hoc analysis. Statistical significance was set at α = 0.05 (*p* < 0.05). Statistical analyses and data visualization were performed in R (version 4.2.3).

## Acknowledgments

This work was funded by NIH Grants R01NS091027 (KMW), R01NS126247 (ES) CureCMD (KMW), F31NS120649 (JNJ), P30NS061800 (OHSU ALM), VA I01-BX004938 (ES), Department of Defense W81XWH-18-1-0598 (ES). The contents of this manuscript do not represent the views of the US Department of Veterans Affairs or the US government.

---

**Supplemental Figure 1.**
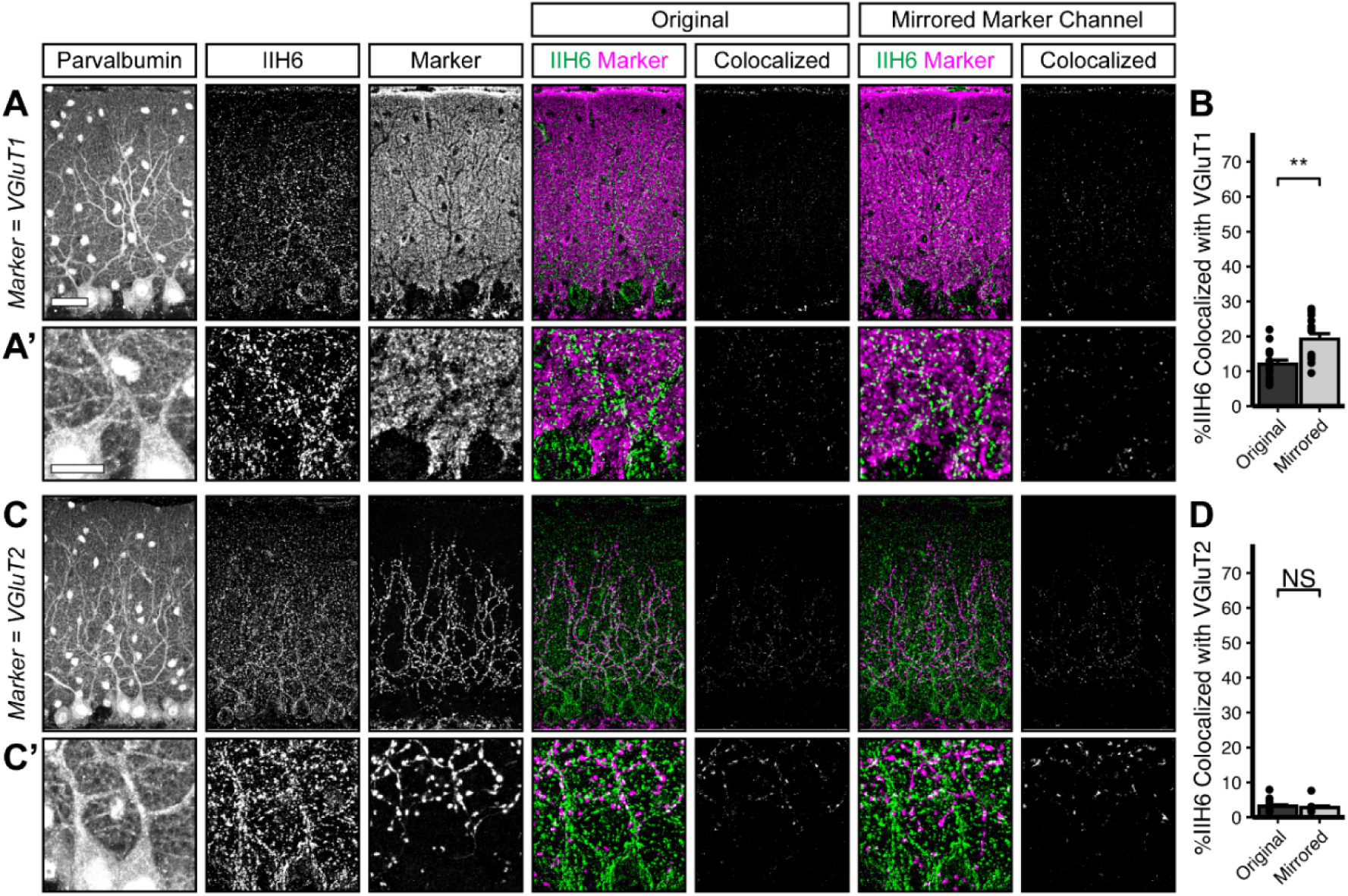
Dystroglycan does not co-localize with markers of excitatory synapses. Cerebellar cortex of lobules V-VI were immunostained with Parvalbumin to show Purkinje cell and MLI morphology and counterstained with IIH6 (glycosylated Dystroglycan) and VGluT1 (parallel fibers) (**A**) or VGluT2 (climbing fibers) (**C**). Both the merged channels (IIH6, green; VGluT1/VGluT2, magenta) and colocalized pixels are shown for the original image and for original IIH6 with the mirrored VGluT1/VGluT2 channel. Images are maximum projections. (**B, D**) Quantification of the percent of IIH6 puncta that are colocalized with VGluT1/VGluT2 puncta. Scale bar for (**A**, **C**) is 50μm; scale bar for insets (**A’**, **C’**) is 25μm. VGluT1 N = 15 images, 3 animals. VGluT2 N = 15 images, 3 animals.

**Supplemental Figure 2.**
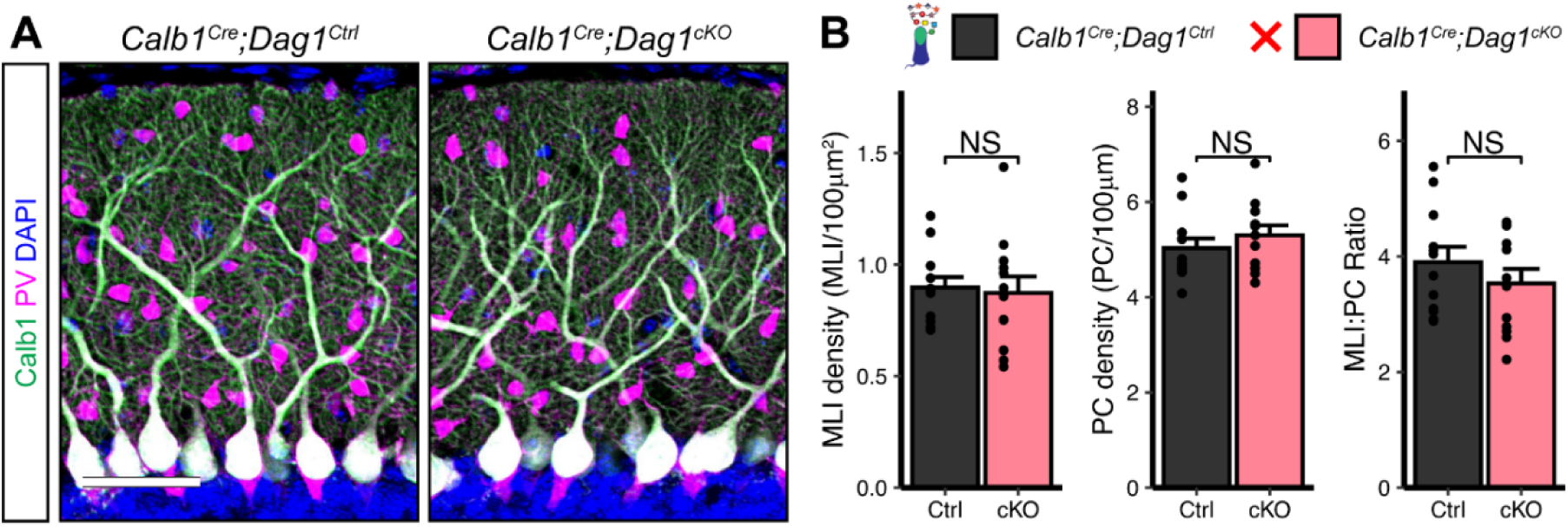
Purkinje and MLI cell counts are unchanged in *Calb1^Cre^;Dag1^cKOs^*. **(A)** *Calb1^Cre^;Dag1^cKO^*and littermate controls immunostained for Calbindin (Purkinje cells, green) and Parvalbumin (Purkinje cells and MLIs, magenta). Nuclei are shown in blue. Images are maximum projections. Scale bar = 50μm. **(B)** Quantification of MLI density, Purkinje cell density, and the ratio of MLIs to Purkinje cells. Error bars represent mean + SEM. *Calb1^Cre^;Dag1^Ctrl^*N = 12 ROIs, 6 images, 3 animals. *Calb1^Cre^;Dag1^cKO^*N = 12 ROIs, 6 images, 3 animals.

**Supplemental Figure 3.**
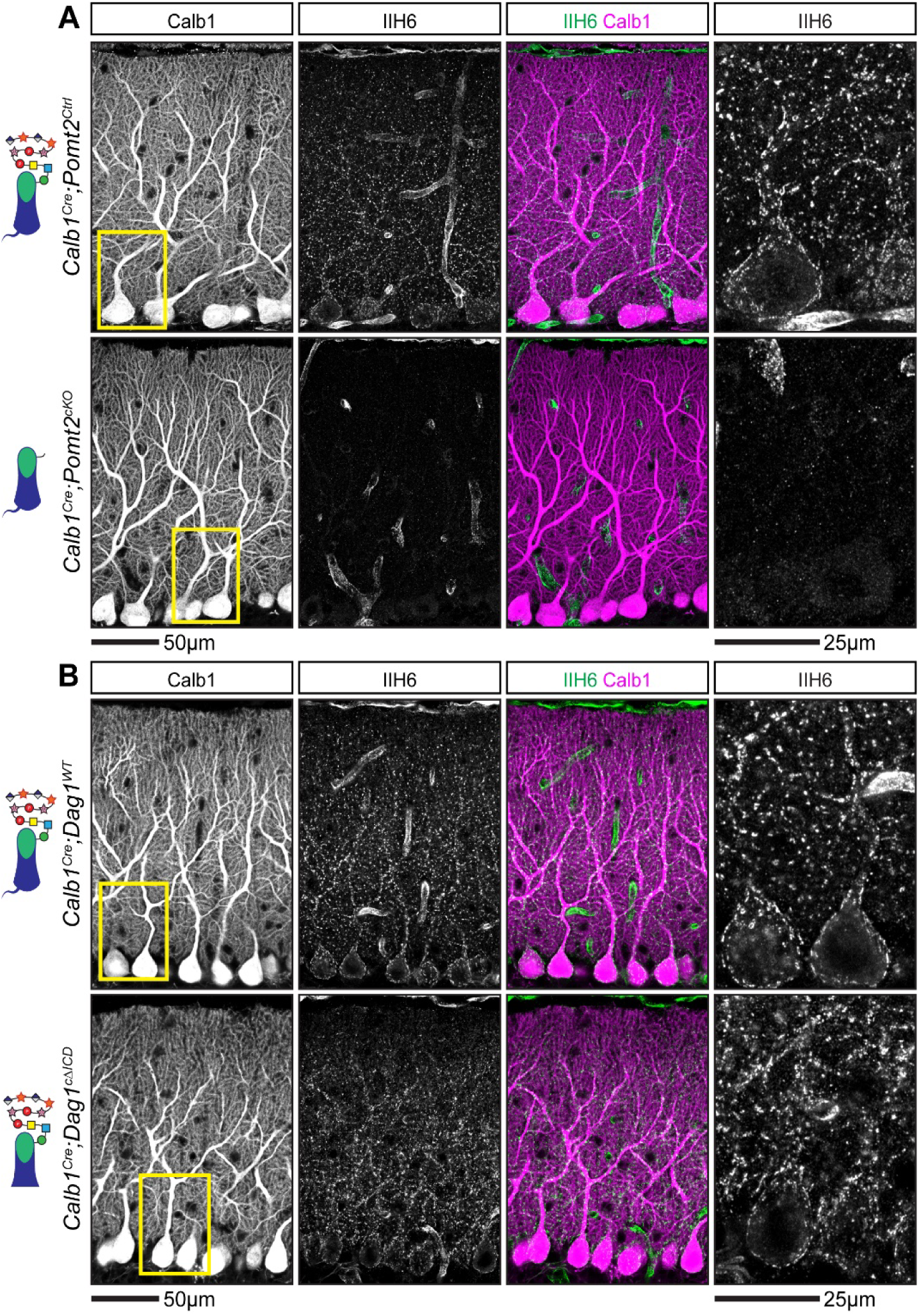
Localization of matriglycan chains on Dystroglycan in *Calb1^Cre^;Pomt2^cKO^* and *Calb1^Cre^;Dag1^c^*^Δ*ICD*^ Purkinje cells. **(A-B)** Cerebellar sections from *Calb1^Cre^;Pomt2^cKOs^* and littermate controls **(A)** or *Calb1^Cre^;Dag1^c^*^Δ*ICDs*^ and littermate controls **(B)** were immunostained for Calbindin (magenta), to visualize Purkinje cells, along with IIH6 (green), to visualize matriglycan chains on Dystroglycan. The rightmost panel represents a magnified view of the area outlined in yellow in the leftmost low magnification panel. Images are maximum projections.

